# Modulation of leg trajectory by transcranial magnetic stimulation during walking

**DOI:** 10.1101/2024.10.23.619051

**Authors:** H. Bourgeois, R. Guay-Hottin, E.-M. Meftah, M. Martinez, M. Bonizzato, D. Barthélemy

**Affiliations:** Institute of Biomedical Engineering, Montréal, QC, Canada; Electrical Engineering, Polytechnique Montréal, Montréal, QC, Canada; Mila, Québec AI Institute, Montréal, QC, Canada; Centre de recherche interdisciplinaire en réadaptation (CRIR), Montréal, QC, Canada; Département de Neurosciences and Centre interdisciplinaire de recherche sur le cerveau et l’apprentissage (CIRCA), Université De Montréal, Montréal, QC, Canada; CIUSSS du Nord-de-l’île-de-Montréal, Qc, Canada; École de Réadaptation, Université De Montréal, Montréal, QC, Canada

**Keywords:** Transcranial Magnetic Stimulation, locomotion, foot clearance, swing phase

## Abstract

The primary motor cortex is involved in initiation and adaptive control of locomotion. However, the role of the motor cortex in controlling gait trajectories remains unclear. In animals, cortical neuromodulation allows for precise control of step height. We hypothesized that a similar control framework applies to humans, whereby cortical stimulation would primarily increase foot elevation.

Transcranial magnetic stimulation (TMS) was applied over the motor cortex to assess the involvement of the corticospinal tract over the limb trajectory during human walking. Eight healthy adults (aged 20-32 years) participated in treadmill walking at 1.5 km/h. TMS was applied over the left motor cortex at an intensity of 120% of the threshold to elicit a dorsiflexion of the right ankle during the swing phase of gait. Electromyographic (EMG) measurements and three-dimensional (3D) lower limb kinematics were collected.

When delivered during the early swing phase, TMS led to a significant increase in the maximum height of the right toe by a mean of 40.7% ± 14.9% (25.6mm ± 9.4 mm, p = 0.0352) and knee height by 57.8%± 16.8%; (32mm ± 9.3 mm; p = 0.008) across participants.

These findings indicate that TMS can influence limb trajectory during walking, highlighting its potential as a tool for studying cortical control of locomotion.

## Introduction

Rhythmic locomotion is predominantly a spinal process, driven by automated movements even in the absence of conscious control (Grillner, 2003). The initiation, modulation and adaptation of walking are controlled by a complex interplay of multisensory feedback, postural reflexes and supraspinal signals (Dietz et al., 2002; Zehr et al., 2004, 2009). Spinal activity during walking can be actively modulated by cortical circuits, which may be responsible for precise muscle control during human locomotion. The importance of the corticospinal tract in locomotion and gait recovery has been extensively studied, particularly using techniques like TMS, a non-invasive technique that involves applying magnetic fields to specific regions of the brain to assess the excitability of cortical and corticospinal pathways (Petersen et al., 2001; Nielsen, 2003; Barthelemy & Nielsen, 2010).

Furthermore, applying repetitive TMS (rTMS) at high frequencies over the motor cortex of patients with incomplete spinal cord injuries resulted in increased excitability in the corticospinal tract (CST) and lead to locomotor improvements, further emphasizing the role of CST in gait recovery(Benito et al., 2012; Krogh et al., 2022).Namely, patients that received rTMS showed significant improvements in the LEMS (Lower Extremity Motor Score), MAS (Modified Ashworth Scale), 10MWT (10-Meter Walk Test), cadence, step length, and TUG (Timed Up and Go), compared to those who received sham stimuli. However, these studies did not include kinematic analyses of gait differences and the impact of corticospinal excitability on gait pattern remains unclear. Recent research indicates that applying TMS during robot-assisted walking leads to a reorganization of spinal reflex pathways (Pulverenti et al., 2022). However, the artificial nature of robot-assisted walking limits kinematic analysis and may restrict the generalizability of findings to natural walking situations.

In animal studies, phasic intracortical neurostimulation has been used to highlight the capacity of motor cortical populations to modulate gait trajectories and coordinate hindlimb movements during locomotion, emphasizing a role in fine-tuning walking patterns (Duguay et al., 2023; Massai et al., 2023). “Phasic” refers to stimuli delivered at precise timings during gait phases. When applied during the swing phase, cortical stimulation has been found to reduce foot-drop, increase flexion capacity, and foster long-term recovery of skilled locomotion capacity in rats with spinal cord injury (Bonizzato & Martinez, 2021).

Building on this previous research, our study aimed at understanding how cortical networks influence the limb trajectory during swing in healthy humans and to determine whether effects similar to those seen in animal models could be obtained. Our main hypothesis posits that stimulation over the leg area of the primary motor cortex will promote increased elevation of the lower limbs during the swing phase of walking. Our secondary hypothesis states that kinematic effects would vary as a function of the stimulus timing within the swing phase.

## Methods

### Aim

Our study aimed at understanding how cortical networks influence the limb trajectory during swing in healthy humans and to determine whether effects similar to those seen in animal models could be obtained. Our main hypothesis posits that stimulation over the leg area of the primary motor cortex will promote increased elevation of the lower limbs during the swing phase of walking. Our secondary hypothesis states that kinematic effects would vary as a function of the stimulus timing within the swing phase.

### Experimental Design

The experimental protocol was performed in two parts: first, the hotspot and motor threshold for TMS was determined while the participants were sitting down. These values were then confirmed during quiet standing. In the second part, TMS was applied at different delays during the swing phase to assess its effect on limb trajectory.

### Participants

Eight healthy adults (5 men, 3 women), aged between 20 and 32 years, underwent screening to identify any prior occurrences of seizures, neurosurgical procedures, and the presence of metal or electronic implants, ensuring eligibility for TMS application. Participants provided informed, written consent for the experimental procedures of this study, which was based on an adaptive trial design approved by the Research ethics committee en réadaptation et en déficience physique of the CIUSSS du Centre-Sud-de-l’île-de-Montréal. This study was conducted in accordance with the central tenets of the Declaration of Helsinki.

### Instrumentation and evaluation

#### Electromyography (EMG)

The EMG signal of the tibialis anterior (TA) and soleus (SOL) muscles was recorded using surface Ag-AgCl electrodes (BlueSensor SP, Ambu A/S, Denmark), 9 mm in diameter and spaced 3 cm apart. After prepping the skin with abrasive tape to ensure it was clean and slightly rough, electrodes were placed on the right and left TA and SOL muscles, selected for their role in generating locomotor activity at the ankle (Fitzpatrick & Day, 2004). The electrodes were placed parallel to the muscle fibers in accordance with SENIAM recommendations (Hermens et al., 2000). A reference electrode was positioned on the right tibial tuberosity. EMG signals were amplified (x1000), bandpass filtered (10-1000 Hz), then digitized and sampled at 2 kHz to a computer using a micro1401 interface (Signal software, Cambridge Electronic Design Ltd, UK).

#### Kinematics

A 10-camera Vicon motion analysis system (Ten Vicon Bonita optical cameras, Oxford Matrices, Oxford, UK, and two additional video cameras) with a sampling rate of 100 Hz was used to capture the participants’ movement during treadmill walking. Prior to data collection, 16 reflective markers (14mm in diameter) were placed on the hip and lower body segments of the participants based on bony landmarks, according to the Plug-in-Gait LowerBody Ai marker set.

#### Temporal parameters of gait

The pressure applied by the feet on the ground was recorded using pressure sensors (Robotshop, Ca) positioned under the toes and heels. The sensors under the heels were positioned to detect the initial foot contact to the ground and determine the initiation of the stance phase. Those under the toes were placed to detect the decrease of weight of the toes at the end of stance, and identify the beginning of the swing phase.

### Part 1-Finding hotspot and intensity of TMS during sitting

TMS was applied over the motor cortex during gait using the BiStim2 stimulator (MagStim, Uk) and a custom-made batwing coil (Jaltron, Us). First, in a sitting position, we placed the coil slightly left of the Cz position, using a 45-degree coil orientation angle with respect to the sagittal direction(Richter et al., 2013). The intensity was gradually increased until a motor evoked potential (MEP) was elicited in the right TA. Surrounding positions were then mapped at the same intensity to determine the optimal position, referred as “the hotspot,” where TMS induced the largest MEP for the same stimulation intensity (Rossini et al., 1994). This hotspot position was then recorded as a target in a neuronavigation system BrainSight (Rogue Research Inc, Ca) to control for stability of the subsequent stimulation. The neuronavigation system uses optical sensors attached to the participant’s head and the stimulation coil. In real-time, these sensors track the participant’s head movements, allowing for the detection of any deviation from the stimulation target. After setting the coil, the threshold for the MEP and for the dorsiflexion of the ankle were established: Participants performed an isometric contraction of the dorsiflexors corresponding to 10% of their perceived maximal effort while stimuli were applied. MEP motor threshold was defined as the minimum stimulus intensity required to elicit a recognizable MEP (over 100μV) in 3 out of 5 stimuli (Liepert et al., 1998). Once this threshold was established, we further increased the intensity to determine the dorsiflexion threshold, when a clear dorsiflexion is detected in 3 out of 5 stimuli. We then used an intensity equivalent to 120% of the movement threshold. The hotspot and intensity of stimulation were validated in the standing position to make sure it induced a visible dorsiflexion of the right leg before moving to the second part of the protocol, when TMS was applied during gait

### Part 2-TMS during gait

The participants walked on a treadmill at a speed of 1.5 km/h. Such a slow pace was chosen to generalize to slower walking speeds exhibited by individuals with spinal cord injury (Barthelemy et al., 2010) and to enable greater stability for TMS application. Initially, each participant walked on the treadmill for 1-2 minutes to become accustomed to the speed. Then, the pressure pattern exerted on the sensor under the toe of the right leg was analyzed on-line to identify the level of activity corresponding to the onset of toe-off, when the load on the toe started to decrease. This point corresponded to the onset of the swing phase and constituted the main trigger to determine the timing of application of the stimulation (Figure 1).

**Figure 1:**
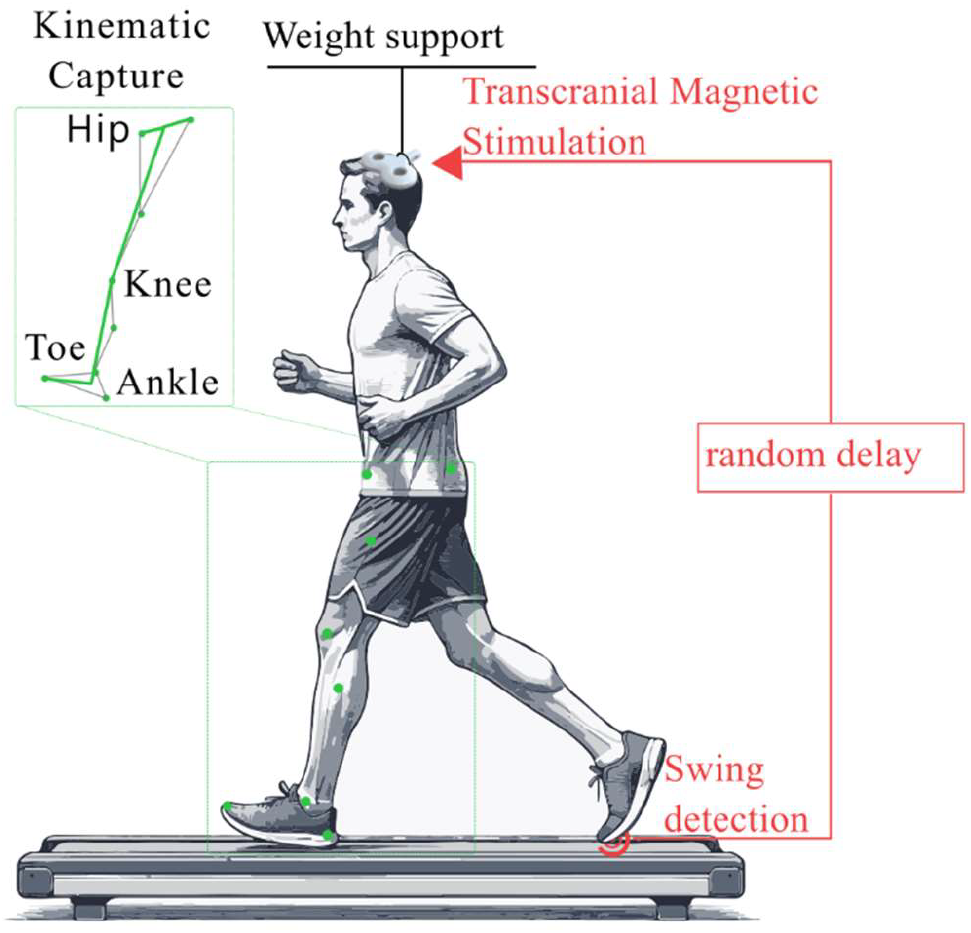
Schematic representation of the experimental setup: As the participant walks on a treadmill, TMS is administered via a coil integrated into a helmet, which is weight-supported. Stimulation is triggered by swing detection through the pressure sensor after a randomized delay. Position markers allow for kinematic tracking.

After the sensors detected the swing initiation, stimuli with a pulse width of 1 ms were administered with delays of 0 ms, 100 ms, 200 ms, 300 ms and 400 ms after toe-off. Twelve stimuli were applied per delay. The order of the stimulation delays was pseudo-randomized, as well as the number of non-stimulated gait cycles between stimulation events, with a minimum of two non-stimulated cycles between each stimulated cycle. The non-stimulated *spontaneous* walking cycles served as a control.

A critical aspect of this setup involved ensuring the stability of the stimulation site amid the rhythmic bodily motions inherent in locomotion while preserving the participant’s comfort and ease of movement. For this purpose, a cyclist’s helmet, modified to provide wide access to the scalp, was utilized. The stimulation coil was mounted on the helmet using adjustable metal fixtures for optimal positioning on the scalp. Due to the heavy weight of the coil (1000 gr) and helmet (665 gr), they were suspended over the participant’s head using a body-weight support system (Anti Gravity Systems LLC, USA). This relieved the weight of the coil and helmet off the participant’s head throughout the entire session (Figure 1). Additionally, the neuronavigation system (BrainSight) was used. If a deviation of 6 mm from the target or more occurred, the experiment was halted, and the helmet position was adjusted to restore the optimal positioning.

### Data analysis

We quantified each MEP as the peak-to-peak measurement of the EMG response volley observed 25 ms after stimulation within a 50 ms interval for each stimulus, and averaged the MEP amplitude for each stimulation delay. To ensure data comparability across participants, MEP amplitudes were normalized relative to each individual’s largest MEP and expressed as a percentage of that maximum MEP. To evaluate the effects of ongoing EMG levels on MEP amplitude, we calculated pre-stimulus EMG activity for each stimulation interval. A pre-stimulus time interval of 50 ms was selected to measure baseline EMG levels, calculated as the average of the rectified signal over the interval. Similarly, to maintain data consistency across participants, EMG activities were normalized relative to each participant’s largest EMG and the results are presented as a percentage of that maximum EMG.

The 3D camera data were processed using Nexus software, followed by custom software, and then MATLAB (R2022b). Toe-off and heel-strike events were identified from pressure sensor traces to segment the data into gait cycles. We analyzed the vertical, frontal, and lateral excursions in the respective vertical, sagittal, and lateral planes of the toe, knee, and hip during each gait cycle. We also examined the angular excursion, which represents the difference between the maximum and minimum angle of the ankle, knee, and hip in all three planes.

Due to variations in swing duration among participants, stimulations with a fixed time-delay did not necessarily coincide with the same sub-period for all participants. Therefore, to ensure consistency, cycles with stimulation were sorted into five bins according to the time when the stimulation occurred within the swing phase. The average duration of the spontaneous swing phase was divided into five equal time bins. Cycles with stimulation occurring in the first 10% of the cycle were sorted in Stim-Bin1, and so forth (Figure 2).

**Figure 2:**
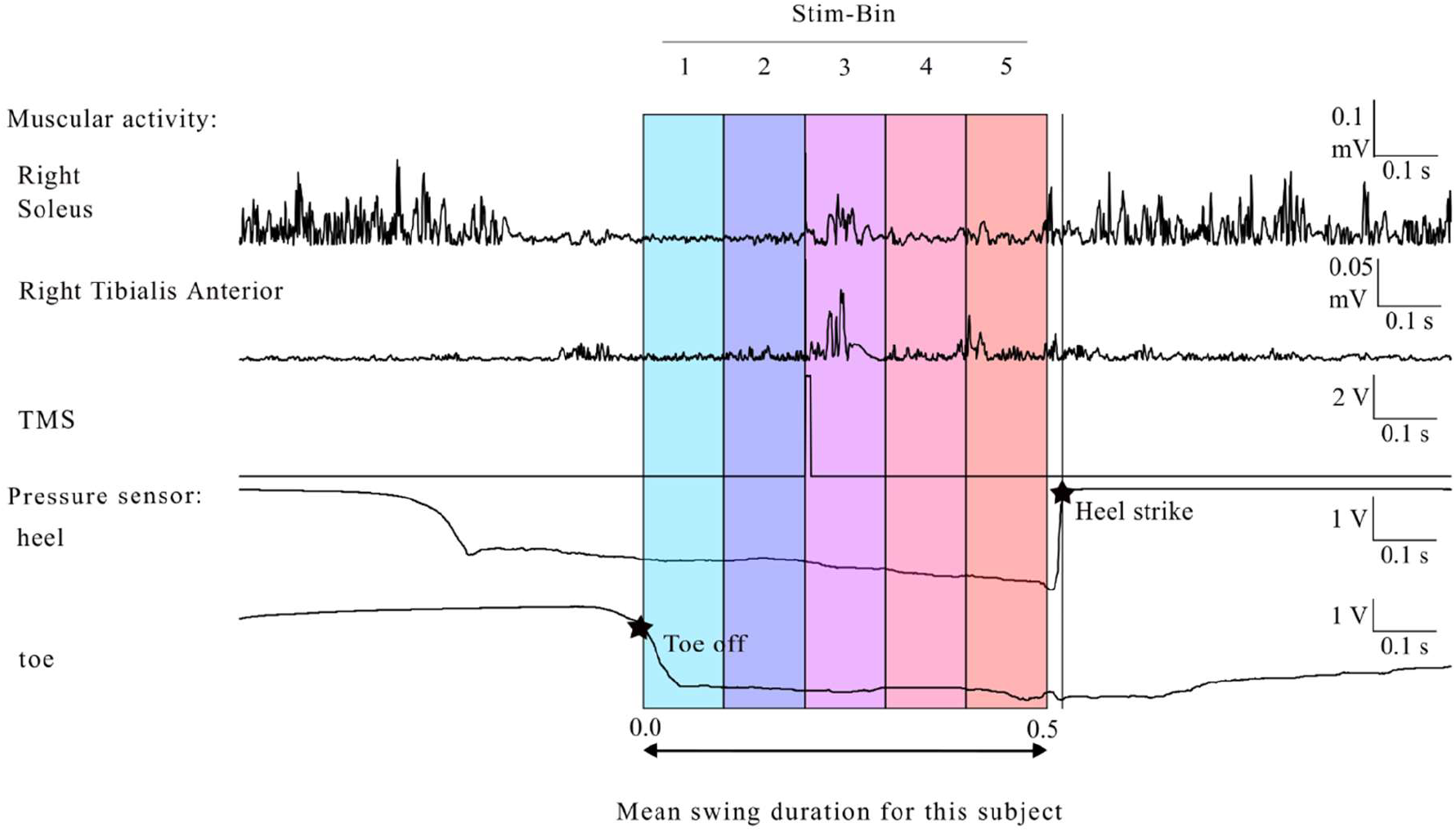
Stimulation labeling based on timing within the swing: Stimulation occurred during the third fifth of the participant’s average swing duration. Consequently, this stimulation is categorized under Stim-Bin3 group. Tracing of the muscle activity of the SOL and TA, TMS stimulation, and the pressure under the heel and toe. The two stars represent toe-off and heel strike.

### Statistical analysis

To determine the differences in kinematic variables between the non-stimulated cycles and the different stim-bins of the stimulated cycles, a one-way repeated measures analysis of variance (RM ANOVA) was conducted. The analysis included eight participants under six different experimental conditions: spontaneous, Stim-Bin1, Stim-Bin2, Stim-Bin3, Stim-Bin4 and Stim-Bin5. Subsequently, a Dunnett post-hoc test was performed to compare spontaneous cycles with each Stim-Bin condition. For the analysis of EMG and MEP data, the same statistical approach was used across the six different experimental conditions: isometric contraction, Stim-Bin1, Stim-Bin2, Stim-Bin3, Stim-Bin4, and Stim-Bin5. Statistical tests were conducted in Graph Pad Prism (v10.2.3). For all statistical tests, significance was set at p< 0.05. Results are presented as mean ± standard error of the mean (S.E.M), reflecting aggregated from different individuals.

## Results

Across all participants, the average number of gait cycles recorded was 219± 11.7, including 44.9± 2.2 stimulated cycle and 169.7±12.5 non-stimulated cycle.

TMS delivered during the early swing modulated the foot trajectory and leg kinematics

Figure 3 illustrates the vertical, lateral and frontal displacement of the toe in one representative participant following stimulation in Stim-Bin1 through 5. An increased vertical toe elevation was observed when TMS was applied in Stim-Bins 1 to 4, with no significant effect observed in Stim-Bin5, corresponding to the latter 20% of the swing phase. A larger lateral displacement was also observed in early swing (Stim-Bin1 and 2), and no significant change was observed in the latter part of swing (Stim-bins 3,4 and 5). Stimulation did not affect frontal displacement.

**Figure 3:**
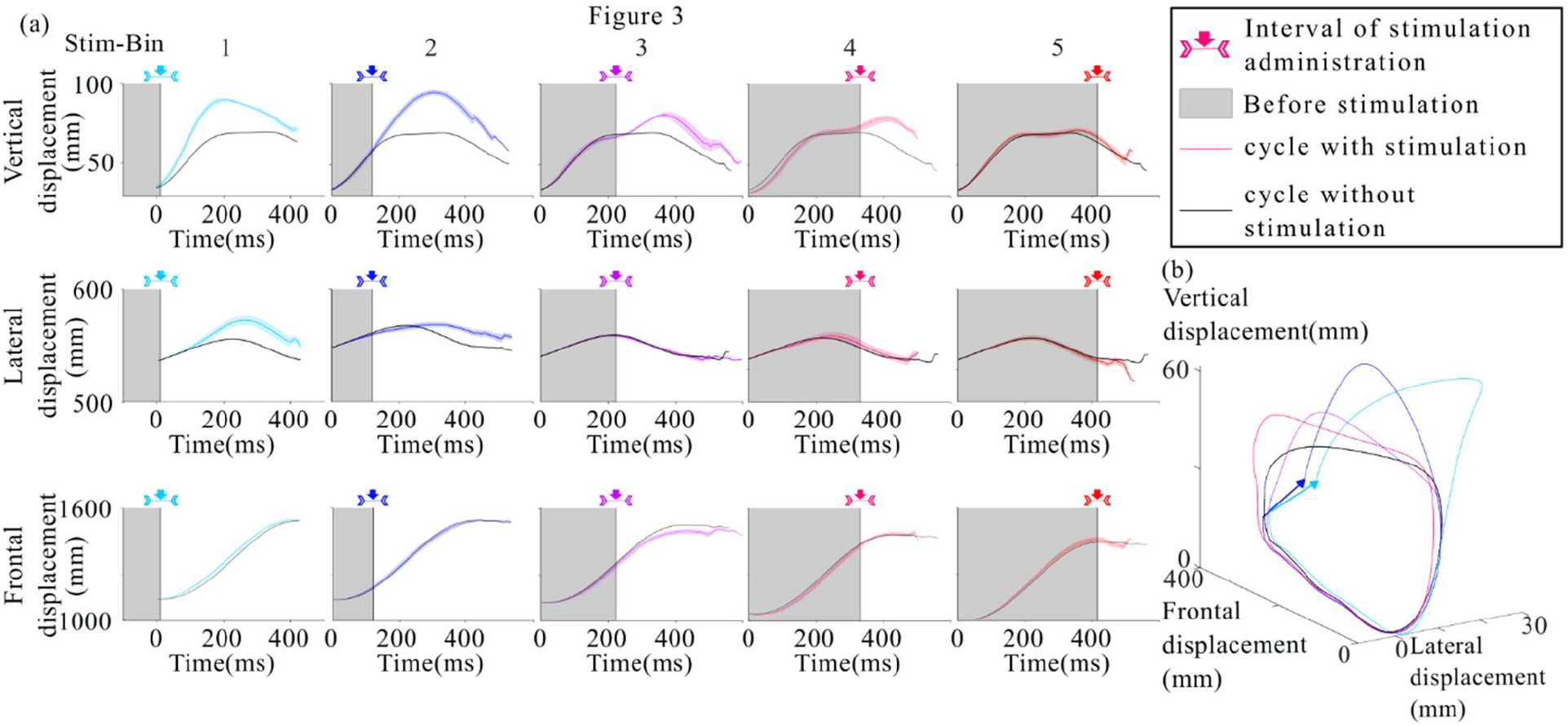
Toe position of a representative participant: (a) Average toe position in the vertical, lateral, and sagittal planes for stimulations in each Stim-Bin compared to the spontaneous cycle, (b) Average toe position in three dimensions for stimulations in each Stim-Bin and the spontaneous cycle. Arrow indicates displacement of the final foot positioning in Stim-Bin 1 and 2.

Similar results were observed across all participants. A significant increase in vertical toe elevation of 40.7%± 14.9 % (25.6mm ± 9.4 mm, p = 0.0352) was observed when stimulation was applied in Stim-Bin1. A similar elevation increase of 43.5%± 17.9 % (27.4mm ± 11.3mm, p = 0.0122) was observed with stimulation in Stim-Bin2, compared to the spontaneous cycle (Figure 4a). Stimulation in Stim-Bin 3, 4 and 5 did not induce significant toe elevation. Lateral toe excursion also showed a significant increase of 4% ± 2.5% (18.1mm ±11mm p= 0.0106) when stimulation was delivered in Stim-Bin1 compared to spontaneous cycles (Figure 4b). No significant differences were observed in the sagittal plane (Figure 4c). The resulting toe trajectories in all three planes are displayed in Fig 4a, b, and c.

**Figure 4:**
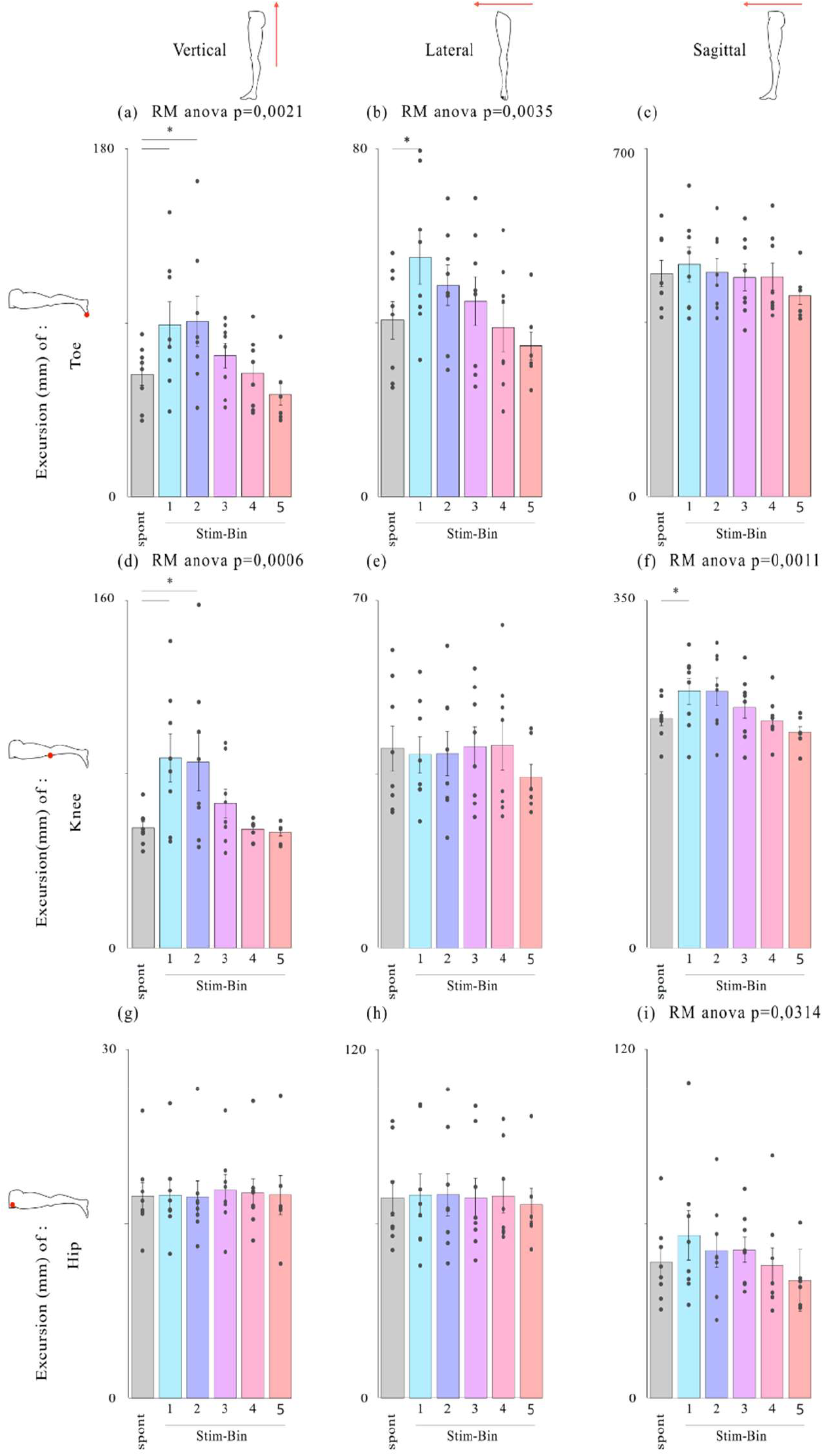
Analysis of leg joint positions for each type of stimulation. Average excursion of the position of the toe(a-b-c), knee (d-e-f) and hip(g-h-i) in the vertical, lateral and sagittal planes, respectively: *p<0.05, Dunnett’s test.

The mean vertical toe excursion in cycles with stimulation consistently exceeded that in cycles without stimulation, although effectiveness varied among participants. Normalized data for all participants are displayed in Fig. 5. This increase in amplitude ranged from 8.58 mm to 73.8 mm (Figure 5a). Lateral and frontal toe displacements showed smaller amplitudes than the vertical displacement but displayed variability across participants, notably participant 2 showing opposite lateral displacement (Figure 5b).

**Figure 5:**
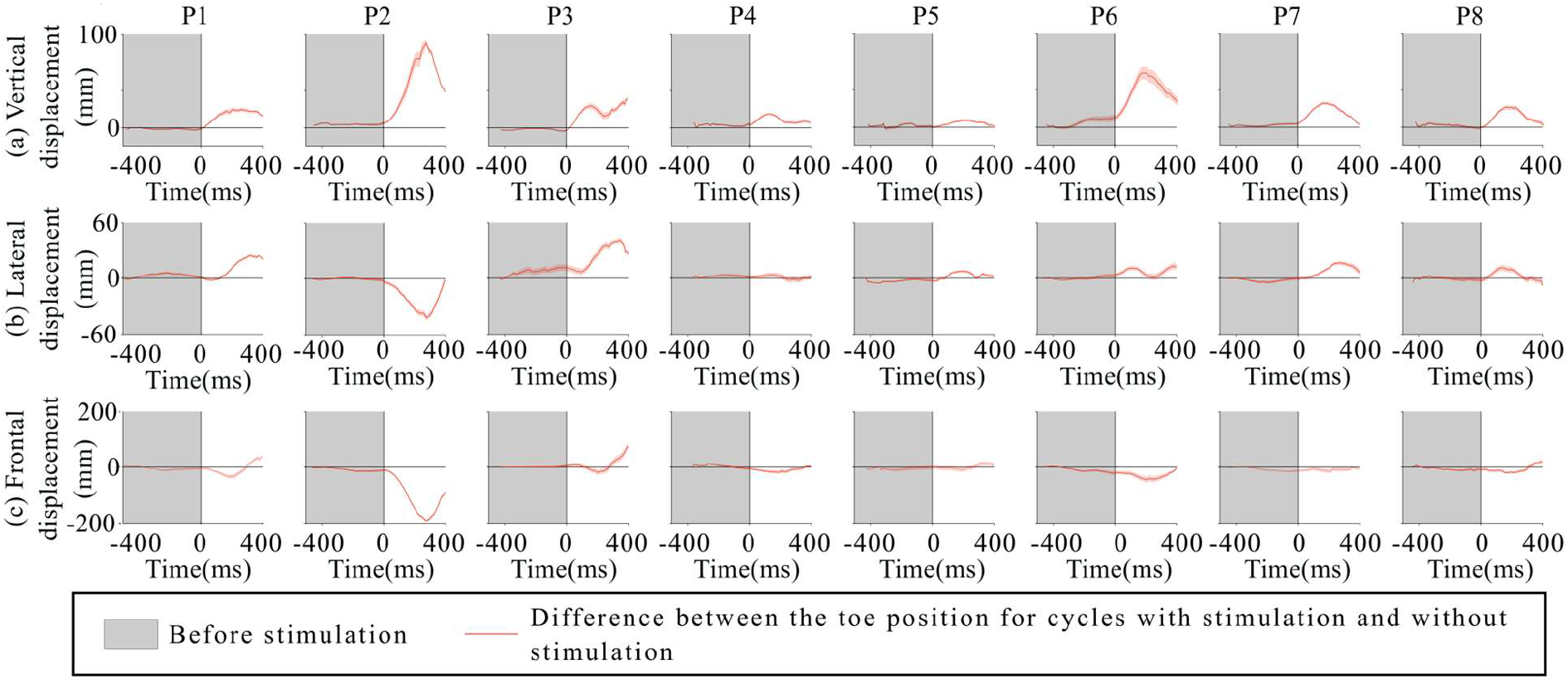
For each participant, the difference between the toe position for cycles with stimulation and without stimulation. (a) in the vertical plane, (b) in the lateral plane, (c) in the sagittal plane. To calculate the displacement, the position of each spontaneous cycle is subtracted from every cycle with stimulation. Thus, the non-shaded portion of the graph represents the average impact of the stimulation immediately after it is delivered.

Although most TMS effects were consistent across participants, individual patterns showed reduced frontal toe excursion in cycles with late stimulation (Stim-Bin3, 4, or 5), particularly noticeable in participants 1, 2, and 6, where stimulation caused premature termination of the swing phase before completing the cycle (Figure 5c).

### Effect of TMS on knee and hip position

Although TMS was optimized to induce activation of the ankle dorsiflexor muscles (namely TA), a significant increase in the vertical knee elevation of 57.8% ± 16.8% (32 mm ± 9.3mm, p= 0,008) was observed when stimulation was applied in Stim-Bin1. This increase was of 54.6% ± 13.2 % (30.2 mm±11.6mm, p = 0,0112) when stimulation was applied in Stim-Bin2, compared to spontaneous cycles (Figure 4d). Furthermore, there was a significant frontal displacement of the knee joint of 11.9% ± 3.1% (27.7mm± 7.3mm, p = 0.0475) when stimulation was delivered in Stim-Bin1 (Figure 4f). No significant change was observed at the knee joint in the lateral plane (Figure 4e) or at the hip in any plane (Figure 4 g, h and i)

### TMS increases ankle and hip angle flexion

Significant increases in angular amplitude during the swing phase (joint flexion, Figure 6) were observed in the sagittal plane for the ankle and hip joints following TMS. Compared to spontaneous cycles, stimulation delivered in early swing resulted in an increase of 36.9% ± 10% in hip angular amplitude (6.5 degree ±1.7degree, p=0.0397, Stim-Bin1, Figure 6c) and an increase of 26.1% ± 6.7 in ankle angular amplitude (4.2 degrees± 1.2 degrees, p=0.0488, Stim-Bin2, Figure 6a). There were no significant increases in angular excursion at the knee. Furthermore, no significant differences were observed in the angular amplitude of the ankle, knee, and hip in other planes (Supplementary feigures 1 and 2).

**Figure 6:**
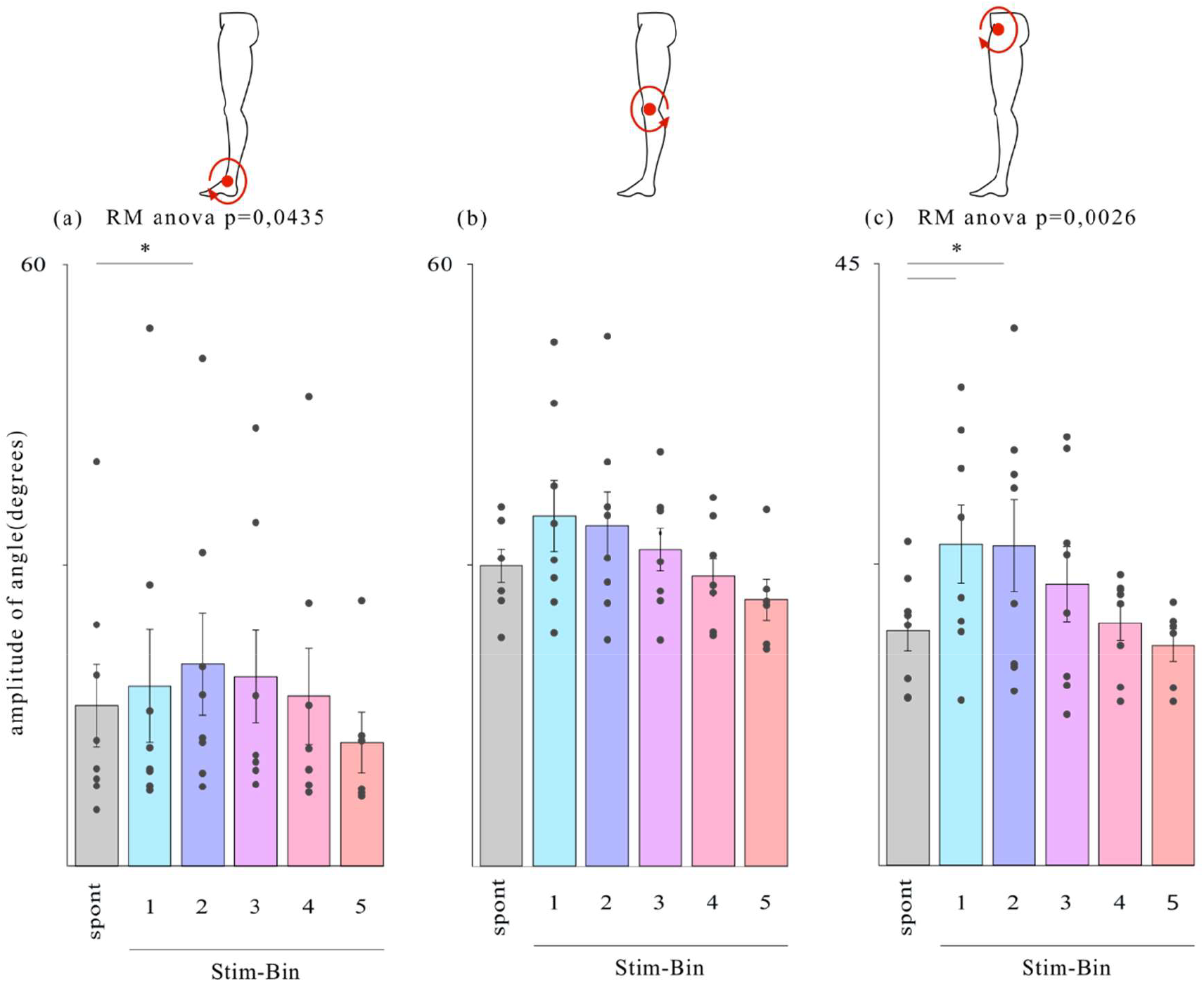
Angular analysis of leg joints for each type of stimulation in the sagittal plane during the swing phase. Average amplitude of the ankle (a), knee (b) and hip (c), *p<0.05, Dunnett’s test.

### Quantification of MEP modulation

To assess changes in corticospinal excitability in the different bis, MEP amplitude increase in MEP amplitudes was observed during each stim-bin of locomotion compared to isometric contraction, both for SOL (p < 0.0001, Figure 7b) and TA (p < 0.0001, Figure 7d). However, no differences were observed between the different stim-bins, contrary to the mean level of TA EMG activity (Figure 7c), which is significantly larger at stim-bin 1 and stim-bin 2. Hence, in both the TA and SOL, the MEP amplitudes remain constant regardless of the timing of stimulation during the swing phase.

**Figure 7:**
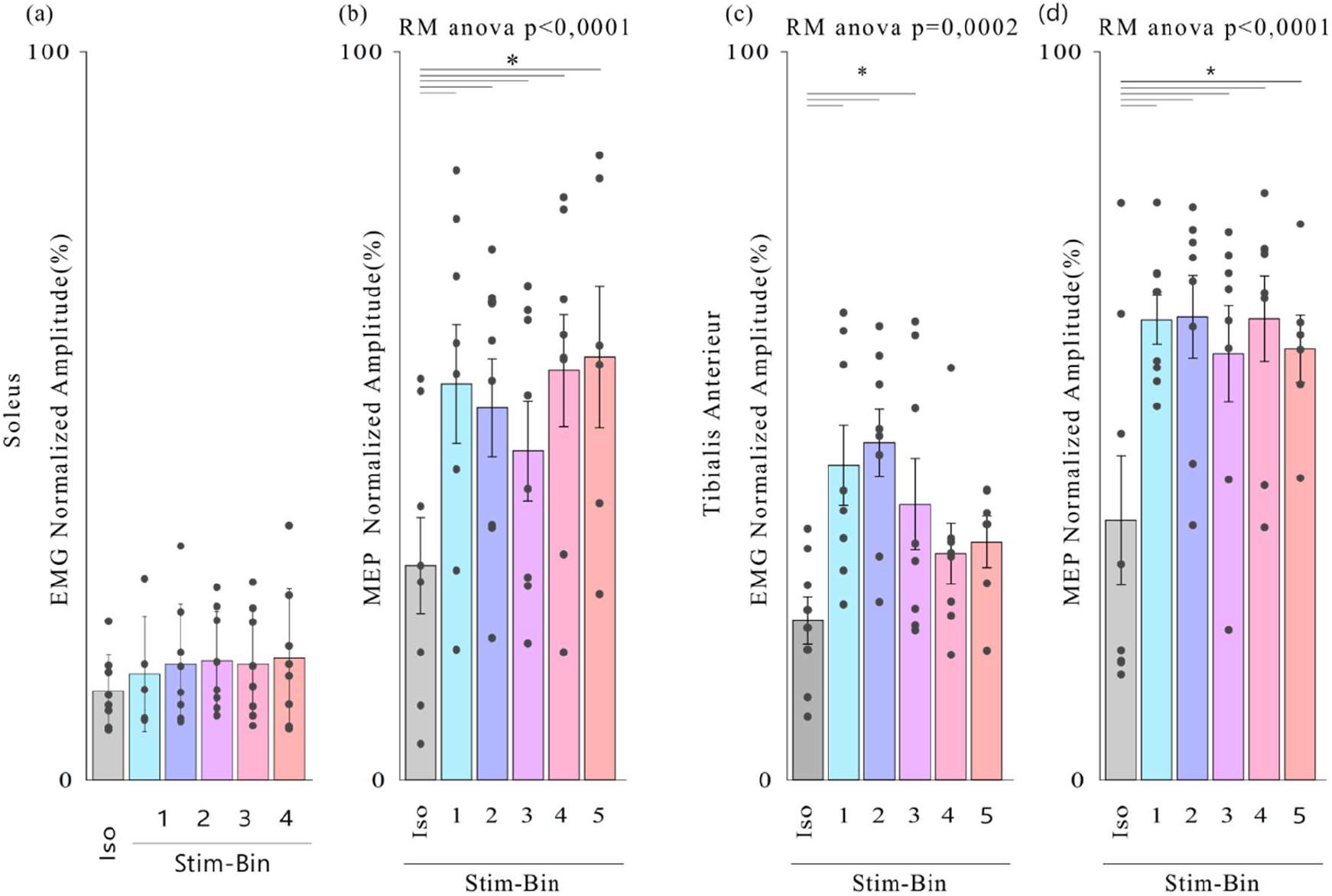
Analysis of muscle activity for each type of stimulation: (a) Background EMG amplitudes before stimulation of the SOL, (b) MEP amplitude of the SOL, (c) Background EMG amplitudes before stimulation of the TA, (d) MEP amplitude of the TA. Iso corresponds to isotonic contraction, *p<0.05, Dunnett’s test.

## Discussion

The results of this study highlight the role of the corticospinal tract in controlling foot trajectory during walking, specifically promoting swing elevation. Results show that TMS increased knee and toe elevation when applied at the beginning of the swing phase, albeit with variability among individuals. TMS also induced movements of the knee and toes in the sagittal and lateral directions when applied at the beginning of the swing phase. MEP amplitudes of the TA and SOL were similar in every stim-bin during gait, regardless of the timing of stimulation during the swing phase or of the mean level of ongoing EMG in TA during gait. As significant changes in trajectory modulation are observed only in Stim-bin 1 and 2, modulation of corticospinal excitability to TA and SOL muscles is not correlated to the changes in the trajectory, and more proximal muscles could be preferably recruited by the stimulation during gait.

### TMS leads to increased foot clearance when applied at the onset of swing

The increase in vertical elevation of the toe during swing may be attributed to increased flexion of the ankle, knee and hip, as measured by angular excursion. Despite tuning stimulation amplitude to achieve suprathreshold effects primarily on ankle dorsiflexion (120% intensity), TMS-induced muscle activation involves the entire leg. The human leg motor cortex is located along the edge and depths of the interhemispheric fissure (Penfield & Boldrey, 1937), where TMS can recruit more superficial cortical networks (Wassermann et al., 2008). Networks of neurons more selectively controlling distal leg movements, particularly the ankle, are located deeper within the fissure, potentially requiring different stimulation techniques for recruitment. The preferential recruitment of flexor muscles echoes previous work done in the cat model; predominance of cortical control over flexor muscles has been extensively shown. Studies (Bretzner & Drew, 2005) showed that stimulation during locomotion evoked larger responses and often activated additional muscles compared to stimulation at rest. Specifically, stimulation during the swing phase increased flexor muscle activity without changing the cycle duration, whereas stimulation during the stance phase decreased extensor muscle activity duration and initiated a new, premature swing phase, thereby resetting the step cycle. Thus, activation of the cortical and corticospinal tracts preferentially recruits flexor muscles, which might be located more proximally. As trajectory is most changed when TMS is allied in early swing, one potential target for TMS might be the semi-tendinous. Indeed, in the cat model, previous studies have recorded discharges from the pyramidal tract neurons within the hindlimb representation of the primary motor (Drew et al., 2002). Results showed that A large proportion of PTNS discharged early in swing, in phase with knee flexors such as the semitendinosus. Others discharged slightly later, in phase with the activity of ankle flexors, such as TA, while still others discharged at the end of swing, in phase with digit dorsiflexors, such as the extensor digitorum brevis. One limitation of the study is that knee flexor or hip flexor muscles were not recorded, which prevents a clear conclusion on this point. Future investigations should look more closely at proximal flexor muscles of the lower limb.

### TMS induces changes in sagittal and lateral trajectories of knee and toe

The induced lateral and frontal changes in the trajectories of knee and toe suggest that TMS affects not only recruit neuronal networks controlling flexion muscles but also neighbouring networks that control slightly different movements. Although these movements were not targeted, they might not be inconsistent with the ongoing walking task. In healthy individuals (Neumann, 2016), an abduction of the ankle and an external rotation of the hip during the swing phase contribute to a lateral external displacement of the toe (Osaki et al., 2007), which was observed in the current study Similarly, in healthy individuals, it is observed during walking that knee flexion during the swing phase results in lateral knee displacement, which was observed in the present study, and is further amplified by TMS stimulation. Invasive microstimulation studies in intact cats have shown that intracortical microstimulation induces various gait patterns, including lateral abductions and diverse foot lifts biased toward the front or back (Duguay et al., 2023). Outputs from neighboring regions can linearly summate (Ethier et al., 2006), suggesting that individual movement outcomes depend on the recruited subregions within the effective cortical activation area, coil positioning, and individual cortical configuration and excitability.

### Constant MEP amplitudes in TA regardless of timing of the stimulation

The consistency of MEP amplitude indicates that the timing of TMS within the gait cycle does not differentially affect corticospinal excitability. This might suggest a stable level of cortical excitability throughout the gait cycle or that MEPs are not sensitive to the specific timing of TMS in this context. This may seem to contrast with previous studies showing strong MEP modulation induced by TMS during walking, likely reflecting changes in cortical excitability (Schubert et al., 1997). Nonetheless, MEP amplitude in TA stayed constant in each stim-bin despite significant trajectory modulation only in the 1^st^ and 2^nd^ stim-bin. This suggests that TA may not be the primary agonist of this trajectory modulation and, as stated above, TMS might preferentially recruit proximal flexor muscles at the hip and knee that can enable this early change in limb trajectory. These results are in line with studies in rats with spinal cord injury : Specifically, neurostimulation applied to the motor cortex effectively lifts the foot during the swing phase of locomotion, involving the entire leg and being most effective when applied early in the swing phase (Bonizzato & Martinez, 2021). Furthermore, in the rat model, the intensity of intracortical neuromodulation correlated with the step height, indicating precise control over leg kinematics. Our observations in humans using TMS as cortical neuromodulation confirm these results and point to similar underlying mechanisms, such as cortical neuronal adaptations, adjustments in neural circuits governing locomotion, and coordinated muscular responses to fine-tune lower limb movements.

### Clinical Implications

TMS is a valuable neurophysiological technique for assessing residual corticospinal connections in patients with spinal cord injuries, thereby facilitating rehabilitation planning. Additionally, TMS evaluates abnormalities in the corticospinal pathway, providing crucial information for tailoring rehabilitation programs and offering promising potential as an adjunct treatment to existing therapeutic protocols. rTMS has been shown to be effective in improving motor function (Jo & Perez, 2020), reducing neuropathic pain(Saleh et al., 2022), and alleviating spasticity(Hou et al., 2020) in patients with spinal cord injuries Unlike conditioning experiments aimed at inducing or reinforcing plasticity, our study introduces an innovative application of TMS: applying it during walking to adjust lower limb trajectory and improve foot elevation. Although technology stimulating at the peripheral nerve already exists to address foot-drop, phase-coherent TMS could thus facilitate flexion at multiple joints and play a beneficial long-term role when integrated into locomotor training, potentially enhancing recovery. This approach offers promising prospects for the development of spinal cord injury rehabilitation therapies, particularly for patients suffering from conditions such as foot drop syndrome during walking, by better understanding the motor control mechanisms of the lower limbs and exploring the potential of TMS to modulate these mechanisms. However, it’s important to highlight that the setup required to administer TMS during walking is cumbersome and would require further engineering to be readily applicable in clinical settings. In particular, the weight support system we used did not fully alleviate the coil weight, which is an important limiting factor when recruiting elder participants or individuals with movement weakness.

Our experimental setup was primarily designed to recruit MEPs in the TA muscle. However, it should be expected that the TA is not the only target. Indeed, angular and EMG activity analyses reveal that knee and hip flexion primarily drive the elevation of the knee and toes. Antagonist SOL was also activated, and stimulation did not result in a significant ankle flexion. This underscores the challenge of precisely targeting the TA muscle with TMS, given the peculiar localization of neurons governing lower limb movements within the motor cortex. Further studies could benefit from recording activity from additional muscle groups to better understand muscle recruitment patterns with TMS (Bonnard et al., 2002; Barthelemy & Nielsen, 2010).

The effects of TMS can also vary in magnitude from one individual to another, however it remained consistently present and reproducible across the entire study cohort. While TMS provides reliable effects, its spatial selectivity is coarse (millimeters to centimeters), compared to local output feature of cortical networks (organized on the scale tens to hundreds of microns, Nudo et al., 1996), suggesting widespread co-activation of several functional subnetwork. An invasive approach with intracortical stimulation was shown to provide more precise control over specific motor outputs during locomotion in the rat (Bonizzato & Martinez, 2021; Massai et al., 2023) and cat model (Duguay et al., 2023), where cortical stimulation primarily modulated foot elevation during swing.

Given the robustness of results across participants and the efficacy in modulating locomotor trajectories, this study may inform the development of further mechanistic neuroscientific studies on cortical control of gait and the development of neuromodulation interventions using either non-invasive or more selective and invasive neuromodulation techniques, potentially beneficial for gait rehabilitation through exercise.

## Conclusion

This study demonstrates that TMS can significantly influence limb trajectory during gait by increasing flexion during the swing phase. TMS most likely recruits neuronal networks controlling flexor muscles located at proximal joints, as no corresponding changes in corticospinal excitability in ankle flexors were observed during gait. These findings shed light on mechanisms enabling the cortex and corticospinal tract to influence the limb trajectory during gait. The findings also suggest that TMS could be used to enhance/correct limb trajectory in individuals with gait impairments. Although the current set-up was cumbersome, these results may lead to the design of other approaches that aim at modifying or enhancing cortical control of gait.

## Supporting information

Data from Figures 4, 6, and 7, as well as from Supplementary Figures 1 and 2.

## Abbreviations

TMS: Transcranial Magnetic Stimulation
EMG: Electromyographic
rTMS: repetitive Transcranial Magnetic Stimulation
MEP: Motor Evoked Potential
TA: Tibialis Anterior
Sol: Soleus
3D: three-dimensional

## Declaration

### Availability of data and materials

All data generated or analysed during this study are included in this published article and its supplementary information files.

## Acknowledgements

We would like to thank the participants for their availability, time, and efforts. We are grateful for the invaluable technical support provided by Y. El Khamlichi, P. Drapeau, and K. Eddaal. Additionally, we appreciate the assistance during the experiments from H. Tonkov.

## Additional Figure

**Additional Figure 1:**
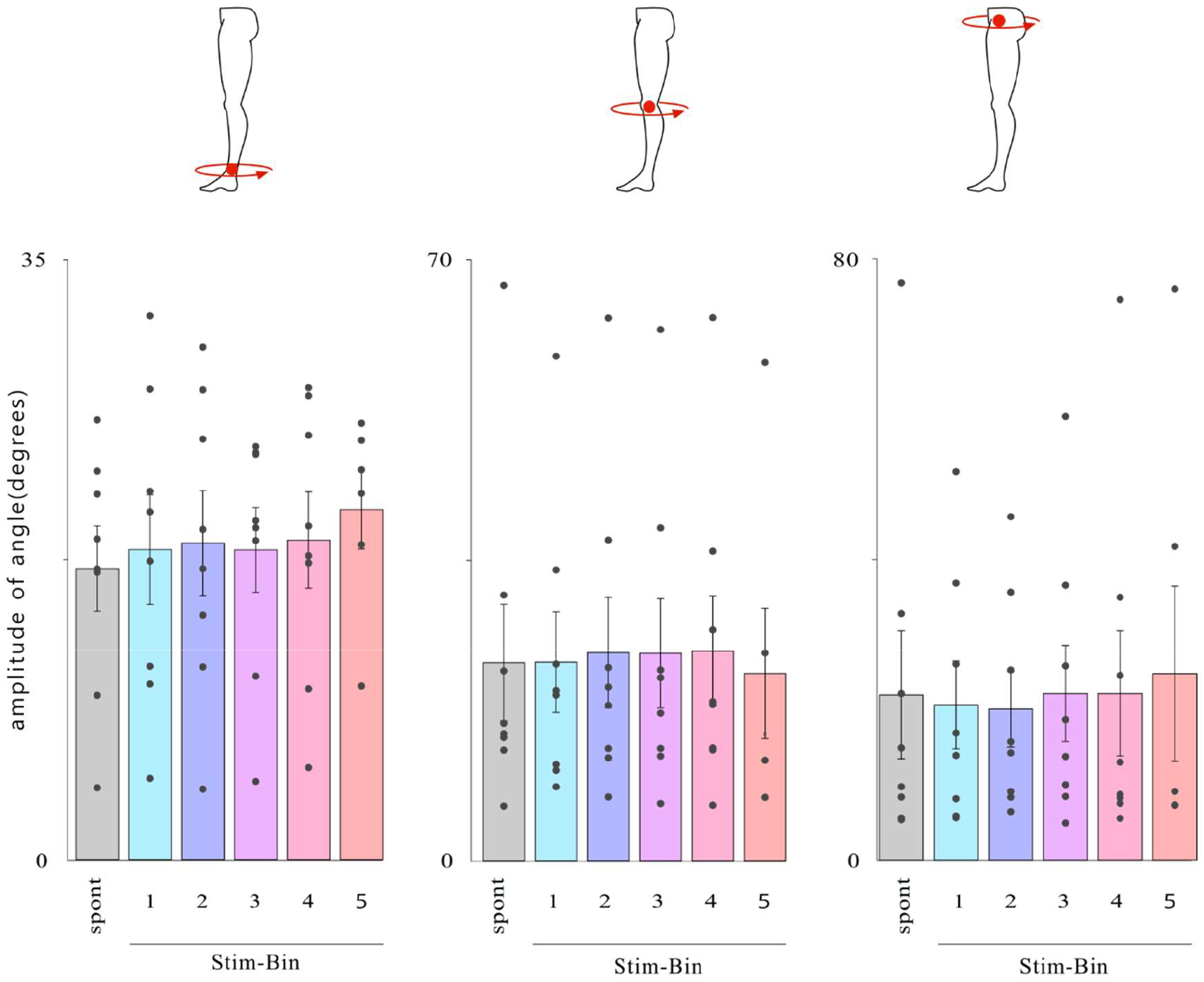
Average excursion during the swing phase of the ankle, knee, and hip angle for each type of stimulation in the lateral plane.

**Additional Figure 2:**
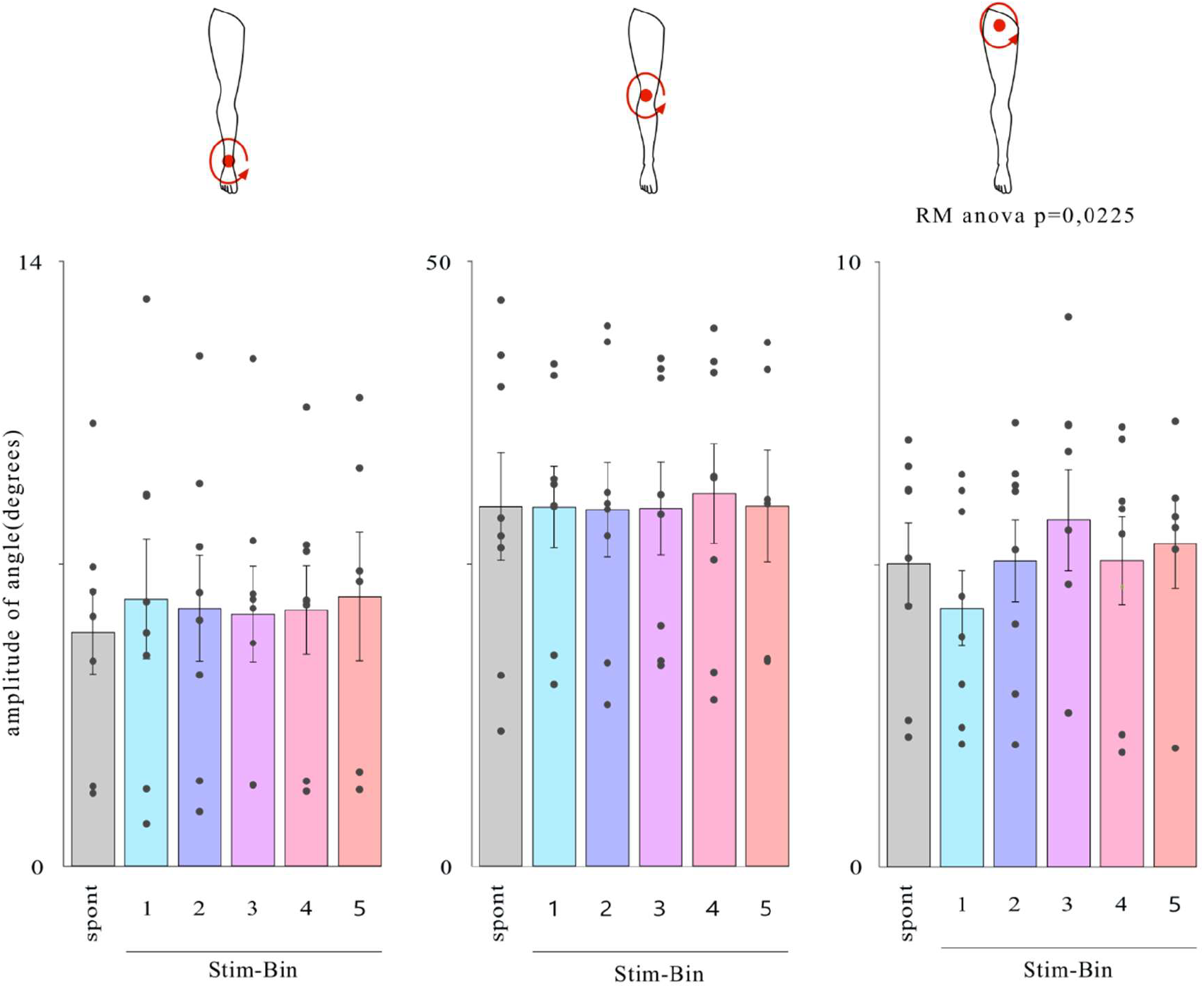
Average excursion during the swing phase of the ankle, knee, and hip angle for each type of stimulation in the vertical plane.

